# Pre-target oculomotor inhibition reflects temporal orienting rather than certainty

**DOI:** 10.1101/2020.05.01.072264

**Authors:** Noam Tal-Perry, Shlomit Yuval-Greenberg

## Abstract

Recent studies suggested that eye movements are linked to temporal predictability. These studies manipulated predictability by setting the cue-target interval (*foreperiod*) to be fixed or random throughout block. Findings showed that pre-target oculomotor behavior was reduced in the fixed relative to the random condition. This effect was interpreted as reflecting the formation of temporal expectation. However, it is unknown whether the effect is driven by target-specific temporal orienting, or rather a result of a more context-dependent state of certainty that participants may experience during blocks with a high predictability rate.

In thus study we dissociated certainty and orienting in a tilt-discrimination task. In each trial, a temporal cue (fixation color change) was followed by a tilted grating-patch. The foreperiod distribution was varied between blocks to be either fully fixed (same foreperiod in 100% of trials), mostly fixed (80% of trials with one foreperiod and 20% with another) or random (five foreperiods in equal probabilities). The two hypotheses led to different prediction models which were tested against the experimental data.

Results were consistent with the orienting hypothesis and inconsistent with the certainty hypothesis, supporting the link between oculomotor inhibition and temporal orienting and its validity as a temporal expectations marker.

## Introduction

Temporal expectation is a prediction regarding the timing of events based on previously-experienced temporal regularities [1]. Recent studies showed that the oculomotor system is tightly linked to temporal attention and expectation: oculomotor activity (saccades, ocular drift, and blinks) is inhibited for a few hundred milliseconds prior to the appearance of a predictable, relative to an unpredictable, stimulus [2–6]. This oculomotor inhibition phenomenon can be used as a sensitive marker for temporal expectation and, as such, has several advantages over traditional behavioral and electrophysiological markers [2]. In our previous studies, we found that oculomotor inhibition prior to a predictable target occurs with various types of temporal structures: with rhythmic expectations, when stimuli were predictable because they are part of a rhythmic stream of stimulation [3], and with associative expectations, when targets are rendered predictable by a precedent informative temporal cue in a visual [2], auditory [5] or tactile [6] task. Oculomotor inhibition was also hypothesized to be modulated by conditional expectations – expectations that gradually increase as time progresses and an expected event has not yet occurred [2]. The oculomotor inhibiton phenomenon is thought to depend on sustained attention as it was reduced in people who were diagnosed with attention deficit and hyperactivity disorder (ADHD) and in participants who showed high inter-trial variability in reaction times, a measurement indicative of sustained attention abilities [3].

In our previous studies on oculomotor inhibition, temporal expectation was manipulated by modulating the foreperiod - the time period between a predicting event (e.g. the onset of a temporal cue or the previous stimulus in a stream) and target onset [2,3,5,6]. In these studies, we included two conditions: (a) a fixed condition, in which the forepriod was identical for all trials, rendering the target predictable and (b) a random condition, in which it varied between a few options, rendering the target less predictable. The findings showed a consistent predictability effect: pre-stimulus oculomotor activity was more inhibited when the foreperiod was fixed relative to when it varied, reflecting the formation of expectations for an upcoming stimulus

Despite the high reliability of the oculomotor inhibition effect and its modulation by target predictability, the characterization of the mechanism underlying this effect is still unknown. Specifically, it is still an open question whether the oculomotor inhibition effect reflects temporal orienting of attention for specific targets. This previously-suggested ‘temporal orienting hypothesis’ is challenged by an alternative ‘certainty hypothesis’, according to which oculomotor inhibition is caused by a general state of elevated certainty in the fixed relative to the random blocks. According to this view, repeated encounters with predictable targets in fixed blocks may have elevated the participants’ general feeling of certainty in those blocks. It could be hypothesized that in states of high certainty, the oculomotor system inhibits activity, either because it is prone to be less explorative or due to lower task engagement.

In the present study we tested the orienting attention hypothesis by contrasting it with the certainty hypothesis and by examining the effects of conditional probabilities. Unlike previous studies that investigated the oculomotor inhibition effects with a variety of oculomotor behaviors (e.g. blinks, drift), in the present study we focus on saccades, which were found to be the most reliable measurement of this effect [2].

In each trial, participants (N=20) were presented with a temporal cue that was followed, after a foreperiod interval, with a target – a tilted grating patch - on which they performed a tilt-discrimination task. Trial-by-trial temporal orienting was modulated by changing the probability of occurrence of specific foreperiods, i.e. certain foreperiods occurred at higher probabilities than others within each block. In contrast, block-wise certainty was manipulated by changing the level of variation between the foreperiods of each block, i.e. in some blocks only a few foreperiods were presented, or even just only, and in other blocks there was a variety of foreperiods. The general state of certainty of the participant within each block, was manipulated by varying the foreperiod distribution between blocks in three different ways: (1) in *full-certainty blocks*, the same foreperiod (1 or 2 s) was used in all trials; (2) in *high-certainty blocks*, one foreperiod (1 or 2 s) was used in most (80%) of the trials and another foreperiod (2 or 1 s, respectively) was used in the rest of the trials; (3) lastly, in *low-certainty block*, five different foreperiods (1, 1.5, 2, 2.5, 3 s) occurred in equal probabilities. These three block types resulted in four trials conditions: (1) *full-certainty trials*: trials of the full-certainty-1s and the full-certainty-2s blocks; (2) *high-certainty-frequent trials*: 1s foreperiod trials of the high-certainty-1s blocks and 2s foreperiod trials of the high-certainty-2s blocks; (3) *high-certainty-rare trials:* 2s foreperiod trials of the high-certainty-1s blocks and 1s foreperiod trials of the high-certainty-2s blocks; (4) *low-certainty trials*: trials of the low-certainty blocks.

By examining the pre-target saccade rate (SR) in 1 s and 2 s foreperiods, we tested two alternative predictions as depicted in **Figure 1**. The *certainty hypothesis* (**Figure 1A**), predicts that the pre-target SR would be determined solely by the foreperiod distribution within the block, regardless of the identity of specific trials. According to this hypothesis, it should be lowest (i.e. strongest oculomotor inhibition) in the full certainty trials, higher in the high certainty trials (both rare and frequent) and highest in the low certainty trials. Importantly, certainty does not vary between foreperiods only between blocks, this hypothesis predicts no significant difference between high-certainty-frequent and high-certainty-rare trials, which occur in the same high certainty blocks. In contrast, the *temporal orienting hypothesis* (**Figure 1B**) predicts that the pre-target SR would be determined exclusively by the probability of a target to occur at a specific timing, regardless of its context within the block. According to this hypothesis, predictions differ for trials with a foreperiod of 1 s and trials with a foreperiod of 2 s. For trials with a 1 s foreperiod, target-probability is determined solely by the frequency of this foreperiod within a given block (20%, 80% or 100%). Accordingly, the hypothesis predicts that pre-target SR should be lowest (i.e. strongest inhibition) in full-certainty trials (in which the target occurs after 1 s in all trials), higher in high-certainty-frequent trials (in which the target occurs after 1 s in 80% of the trials) and highest in high-certainty-rare and the low-certainty trials (in which the target occurs after 1s in 20% of the trials). Importantly, for 1s foreperiod trials the hypothesis predicts no significant difference between high-certainty-rare and low-certainty trials, as the probability for a 1 s foreperiod in both conditions is 20%. For trials with a 2 s foreperiod, on the other hand, temporal orienting is not only determined by foreperiod frequency but also by the conditional probability – how likely a target is to occur at a certain time given that it has not occurred up until that point (the hazard-rate function). Considering that pre-target SR is measured slightly before 2 s post cue, at this time range the probability of target occurrence is full in three out of the four trial-types. Naturally, probability for target occurrence is full in full-certainty trials in which the targets occur at 2 s in all of the trials; but it is also full in the high-certainty trials, where there are only trials with foreperiods of 1 s or 2 s, and therefore once one second has elapsed and no target has yet appeared, the target will certainly occur at exactly 2 s post cue. In contrast, in low-certainty blocks, where there are five optional foreperiods, slightly before 2 s there is only a 33% chance for the target to occur at 2 s. The temporal orienting hypothesis predicts pre-target SR would reflect these conditional probabilities, i.e. be high for the low-certainty blocks and low for the other three block types. Notably, the hypothesis predicts no significant difference between the full-certainty, high-certainty-rare and high-certainty-frequent trials as in all three conditions the target is fully predictable at 2 s, given that it has not occurred at 1 s.

**Figure 1.**
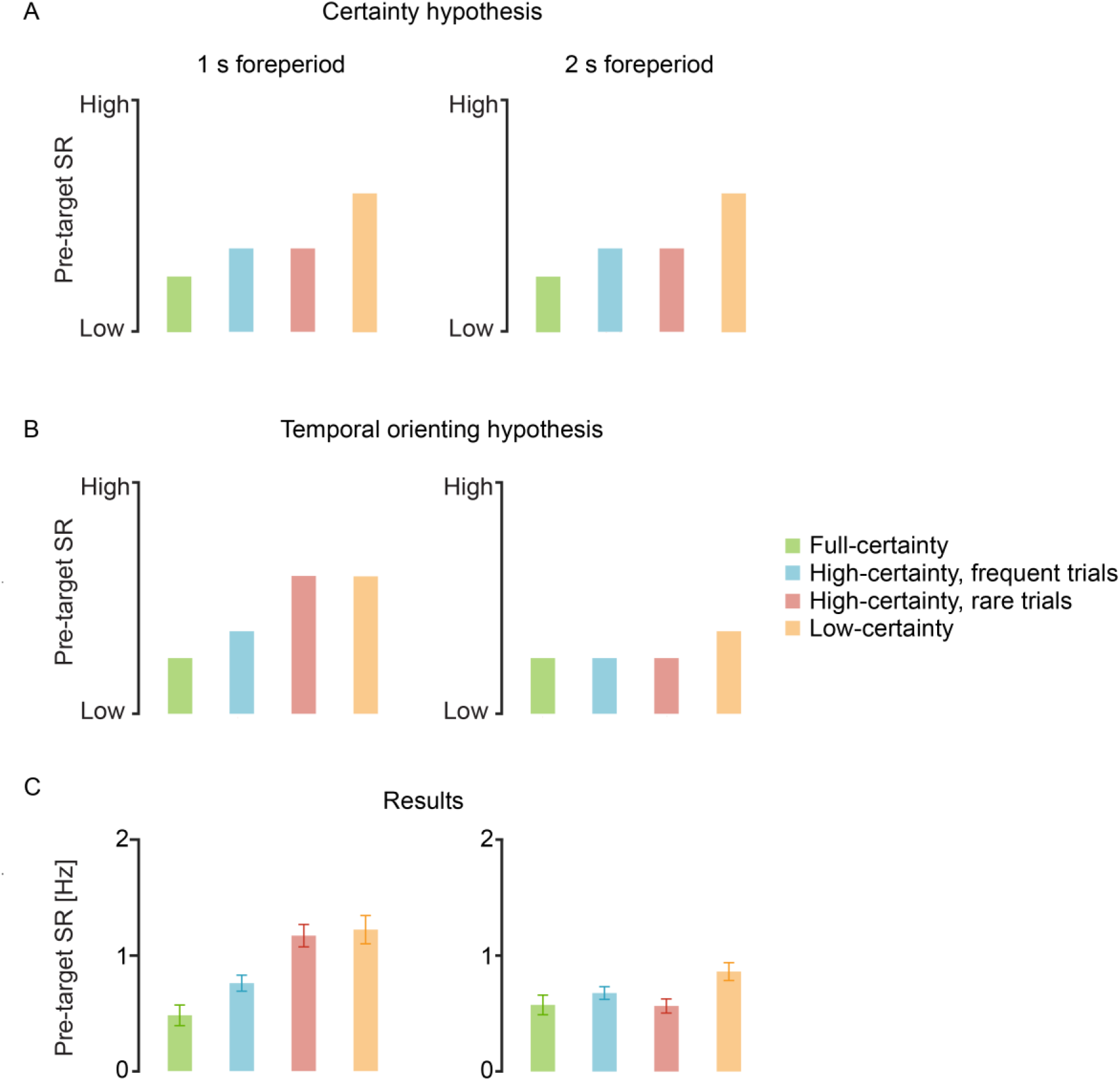
Hypotheses and results. Theoretical predictions of the *certainty* (A) and the *temporal orienting* (B) *hypotheses* for pre-target saccade-rates (SR) in trials of 1 s (left) and 2 s (right) foreperiod, of the four conditions. (C) Grand average pre-target SR of 20 participants, averaged across the time interval of −100 to 0 ms relative to target onset. Error bars designate ± 1 within-subject standard error from the mean [31]

Results of the present study (**Figure 1c**) confirm the predictions of the temporal orienting hypothesis and are inconsistent with the certainty hypothesis. These findings provide conclusive evidence that oculomotor inhibition reflects temporal orienting for specific targets rather than a general sense of certainty, and further validate the use of oculomotor inhibition as a marker of target-specific temporal expectation.

## Methods

### Participants

Twenty students of Tel-Aviv University participated in this experiment (11 female, one left-handed, Mean age 22.6, standard deviation (SD) age 3.5). The sample size was determined based on a similar previous study which used the same number of participants [2]. To ensure that that this sample size leads to adequate power in this study despite having a lower number of trials per condition (80 here compared to 100 in the previous study), we performed a power simulation (using Superpower package [7]) based on truncated data of the previous study using only the 80 first trials of each condition. Simulation results indicated that with 20 participants and 80 trials there is a high probability for finding predictability effects (fixed vs. random) with 1 s (1 − *β* > .99, Cohen’s *d_z_* = 1.902) and 2 s (1 − *β* > .99, Cohen’s *d_z_* = 1.275) foreperiods. Participants reported normal or corrected-to-normal vision with no history of neurological disorders. Participants volunteered or received payment or course credits for their participation. The experiment was approved by the ethical committees of Tel-Aviv University and the School of Psychological Sciences at Tel-Aviv University, and all methods were carried out in accordance with the guidelines and regulations set by the Declaration of Helsinki. All participants signed an informed consent prior to their participation.

### Stimuli

Fixation-target was a cross (black or blue, 0.4° × 0.4°), and target was a Gabor grating patch (2° diameter, 30% contrast, spatial frequency of 5 cycles/degree) slightly tilted clockwise (CW) or counter-clockwise (CCW) from vertical, with tilt degree determined individually via 3-up 1-down staircase procedure (mean 1.02°, SD 0.41°, range 0.5-1.8°). All stimuli were displayed at screen-center on a mid-gray background.

### Procedure

Participants sat in a dimly lit room at a distance of 100 cm from the monitor (24” LCD ASUS VG248QE, 1,920 × 1,080 pixels resolution, 120 Hz refresh rate, mid-gray luminance of 110 cd/m^2^) using head- and chin-rest. The experiment was generated and controlled using MATLAB R2015a (Mathworks, USA) with Psychophysics Toolbox v3 [8]. Trial procedure was similar to a previously reported experiment [2], and is depicted in **Figure 2**. A central black fixation cross was presented between trials for a jittered inter-trial interval of 700-1200 ms. At trial onset, the fixation cross changed color from black to blue. This color change acted as a cue which marked the onset of the foreperiod interval. After the foreperiod had elapsed, the target (tilted Gabor patch) was briefly (33 ms) presented and followed by a blank screen. Participants were asked to report the orientation of the Gabor patch (CW/CCW) by pressing one of two keys. When a response was detected or 3 s with no response had elapsed, the fixation changed color again to provide feedback on task performance (green or red for correct and incorrect answers, respectively). A lack of response was considered as an incorrect response.

**Figure 2.**
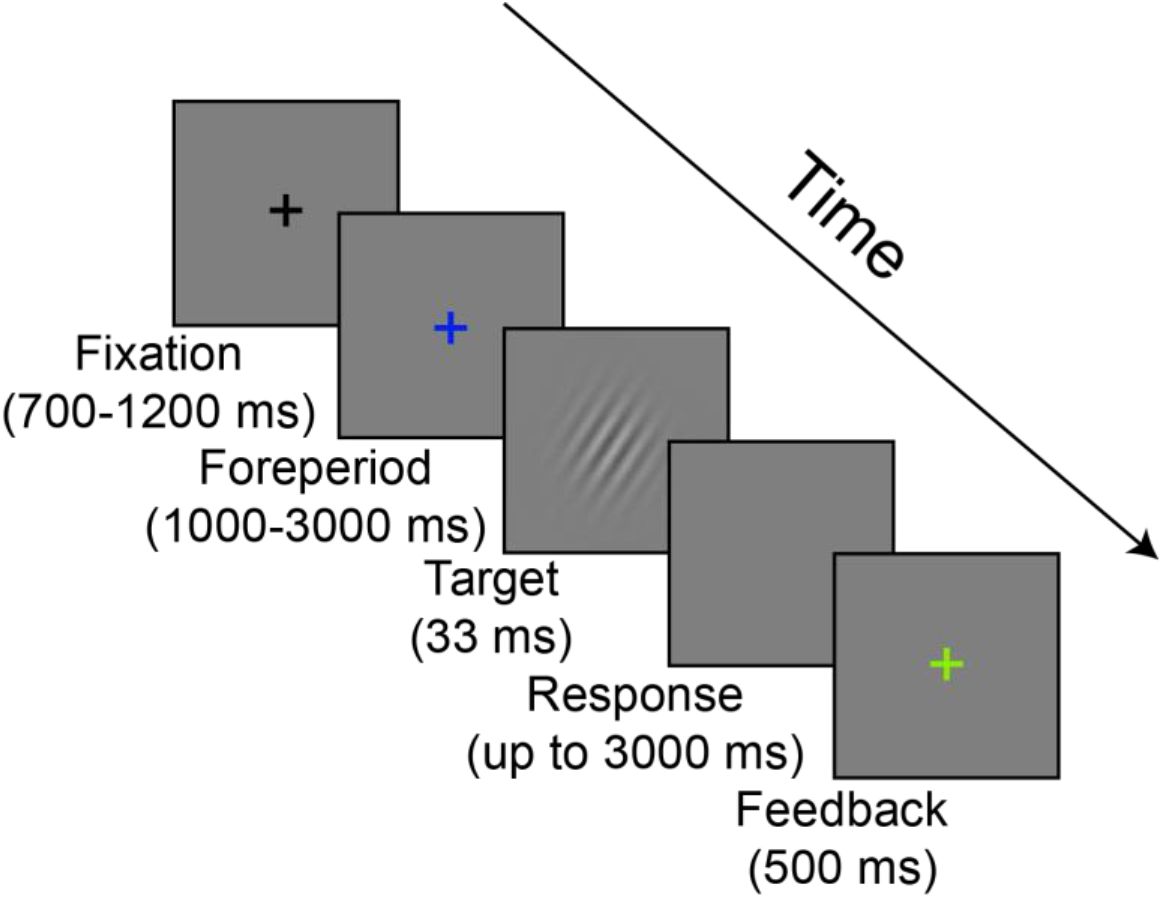
Trial progression. Exemplary trial with a correct response. In incorrect response trials or in trials where no response was given within 3000 ms, the feedback was colored red. The foreperiod prior to target onset is determined according to condition, as depicted in Table 1. Stimuli are shown for illustration purposes only and are not to scale.

**Table 1.** Foreperiod distribution across conditions. Percentage of trials of each foreperiod, within a block.

|  |  | *Foreperiod [ms]* |  |  |  |  |
| --- | --- | --- | --- | --- | --- | --- |
|  |  | *1000* | *1500* | *2000* | *2500* | *3000* |
| *Condition* | *Full-certainty-1s* | 100% | 0% | 0% | 0% | 0% |
|  | *Full-certainty-2s* | 0% | 0% | 100% | 0% | 0% |
|  | *High-certainty-1s* | 80% | 0% | 20% | 0% | 0% |
|  | *High-certainty-2s* | 20% | 0% | 80% | 0% | 0% |
|  | *Low-certainty* | 20% | 20% | 20% | 20% | 20% |

There were five different block types with 80 trials in each block. These block types differed by the distribution of their foreperiod durations, which determined the level of certainty: (1) Full-certainty-1s: a 1s foreperiod was used in all trials; (2) Full-certainty-2s: a 2 s foreperiod was used in all trials; (3) High-certainty-1s: a 1 s foreperiod was used in 80% of the trials and a 2 s foreperiod was used in the rest of the trials; (4) High-certainty-2s: a 2 s foreperiod was used in 80% of the trials and a 1 s foreperiod was used in the rest of the trials. (5) Low-certainty: The foreperiods were uniformly distributed between 1 s to 3 s in 0.5 s increments within each block. Foreperiod distribution for each condition is summarized in **Table 1**. Participants were not informed of the differences between blocks, such that any learning that occurred was incidental.

Participants completed one block of Full-certainty-1s, one block of Full-certainty-2s and five blocks of each of the other block-types. The study consisted of two separate sessions, one with 10 blocks and the other with 7 blocks. All blocks of the same type were performed serially in order to support the learning of the specific foreperiod distribution of that block-type, but the order of the block types within a session was randomized between participants. A short break was given every other block as well as upon the participants’ request. A practice session consisting of 12 trials with a fixed foreperiod of 2 s was administered at the beginning each experimental session. In the first session, a staircase procedure was performed on the Gabor tilt (with fixed foreperiods of 2 s) to determine individual thresholds, yielding accuracy rates of approximately 79%. This threshold tilt was then applied to the rest of the experimental blocks of both sessions. The starting tilt of the staircase procedure was set to 15° and step-size changed logarithmically. The staircase procedure was terminated after four consecutive reversals, or after reaching 100 trials.

### Eye tracking

Binocular eye movements were monitored using EyeLink 1000 Plus infrared video-oculographic desktop mounted system (SR Research Ltd., Oakville, ON, Canada), which has a spatial resolution lower than 0.01° and 0.25–0.5° average accuracy when using a chin-rest, according to the manufacturer. Eye-gaze position was recorded continuously in each block at a sampling rate of 1000 Hz. A nine-point calibration was performed at the beginning of the experiment as well as when necessary.

Gaze analysis was done using MATLAB R2018b. The raw gaze data was low-pass filtered at 60 Hz (as in [2]), and segmented between −300 ms relative to cue onset and 300 ms relative to target onset. Blinks were detected based on the built-in algorithm provided by EyeLink, employing an additional criterion requiring binocular change in pupil size that exceeded 2.5 standard deviations from segment’s mean pupil-size for 3 or more consecutive samples [9].

Saccades were detected using a published algorithm [10,11], such that saccade onset was defined as the point in which the absolute standardized eye velocity exceeded the segment’s median eye velocity by six or more standard deviations, for a minimum of six consecutive samples. Only binocular saccades were included in the analysis. A 50 ms interval between saccade offset and next the saccade onset was imposed to prevent detection of overshoots [12]. Intervals with blinks and 200 ms before the onset of blinks and after their offsets, were excluded from the saccades analysis. Analysis included saccades of all sizes, although most saccades were miniscule (<1°) due to the instruction to maintain fixation. Correlation between saccade amplitude and peak velocity (main sequence [13]) was high (r > 0.9) for all participants.

For each participant and condition, saccade-rate (SR) per second was calculated by averaging the number of saccade onsets per sample across all samples (not including samples that were excluded due to blinks), and multiplying these values by the sampling rate (1000 Hz). Following previous studies [2,3], statistical analysis was performed on the *mean pre-target SR* measurement: average saccade rate at −100 to 0 ms relative to target onset.

### Statistical analysis

There were four trials conditions: (1) *full-certainty;* (2) *high-certainty-frequent;* (3) *high-certainty-rare;* and (4) *low-certainty*. Mean pre-target SR was analyzed using a two-way repeated measures ANOVA, with Condition (full-certainty, high-certainty-frequent, high-certainty-rare and low-certainty) and Foreperiod (1 and 2 s) as independent variables. This ANOVA was followed by simple effect analysis on the two foreperiods, using planned contrasts with False-Discovery-Rate (FDR) adjusted p-value [14]. Sphericity was tested using Mauchly’s test, and corrected *p*-values are reported for significant violations (*p* < .05) along with epsilon (*ϵ*) value, with correction method determined according to epsilon (Greenhouse-Geisser for (*ϵ* < .7; Huynh-Feldt for (*ϵ* > .7). Within-subject standardized effect-sizes (Cohen’s *d* or 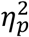, according to analysis) are reported for each significant (*p* < .05) result along with the 95% confidence interval (CI).

Support for null effects is provided by reporting the Bayes factor (BF) for non-significant results (*p* > .05) along with approximate percentage error of the estimates. BFs were calculated using a default (standard) Cauchy prior for the alternative hypothesis [15]. For convenience, BFs are reported with the null hypothesis in the nominator (*BF*_01_), where the reported value indicates how many times the results are more likely under the null hypothesis, compared to the alternative hypothesis. Analyses were conducted using R-studio v1.1.463 [16] in R v3.5.2, with afex package [17] used to perform analysis of variance, with emmeans [18] and sjstats [19] packages used to calculate effect sizes. BF was calculated using the BayesFactor package [20].

An additional analysis aimed at examining the effect of conditional probabilities on saccade rate was conducted on all five foreperiods of the low-certainty condition and reported in Supplementary Material S1. This analysis revealed a replication of the previous reported [2] ‘foreperiod effect’ of saccade rate – a reduction on pre-target saccade rate when foreperiods become longer. We discuss this finding in detail in Supplementary Material S1.

## Results

### Behavior

Accuracy rates and reactions times are reported in **Tables 2** and **3**, respectively. Since there were no specific hypotheses regarding these measures, no statistical tests were performed on them.

**Table 2.**
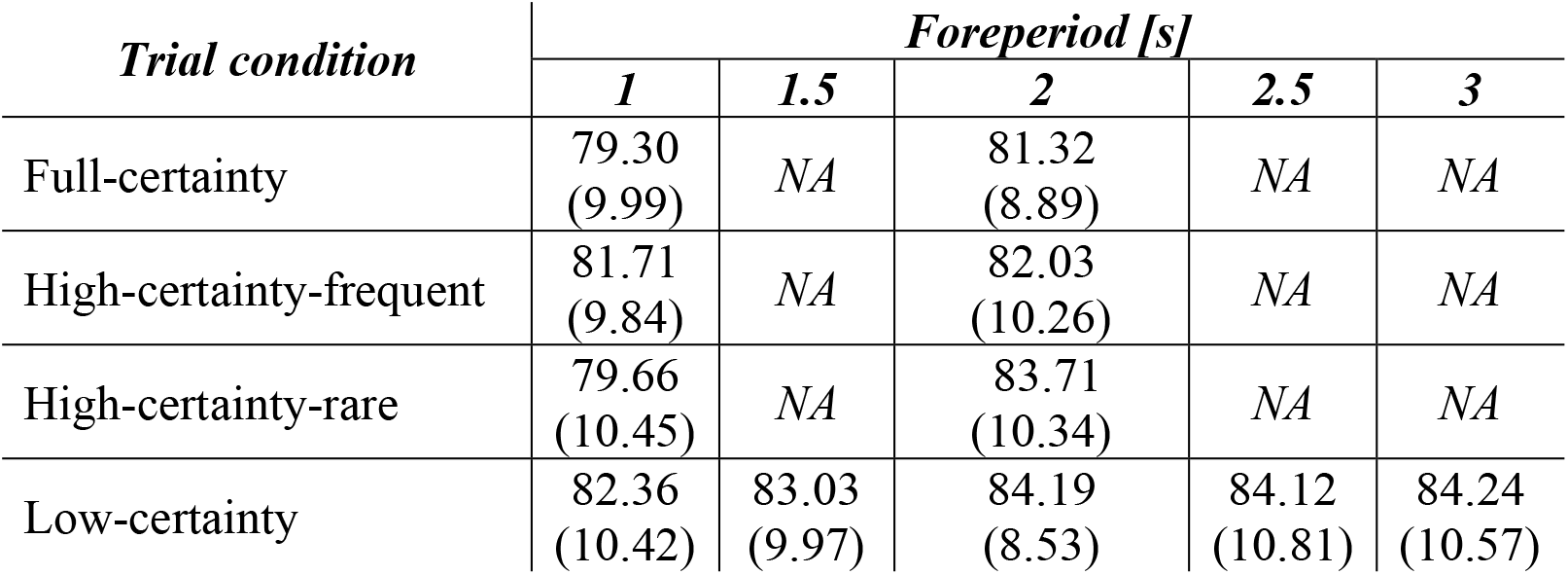
Mean accuracy [%], averaged on the data of 20 participants separately for each condition and foreperiod. Standard deviations [%] are reported in parentheses. Trial conditions with no corresponding foreperiod marked as non-applicable (NA).

**Table 3.** Mean reaction times for correct trials [ms], averaged on the data of 20 participants separately for each condition and foreperiod. Standard deviations [ms] are reported in parentheses. Trial conditions with no corresponding foreperiod marked as non-applicable (NA).

| ***Trial condition*** | ***Foreperiod [s]*** |  |  |  |  |
| --- | --- | --- | --- | --- | --- |
|  | ***1*** | ***1.5*** | ***2*** | ***2.5*** | ***3*** |
| Full-certainty | 648<br>(168) | NA | 657<br>(149) | NA | NA |
| High-certainty-frequent | 665<br>(174) | NA | 651<br>(126) | NA | NA |
| High-certainty-rare | 681<br>(148) | NA | 664<br>(165) | NA | NA |
| Low-certainty | 661<br>(112) | 634<br>(100) | 640<br>(110) | 626<br>(89) | 640<br>(96) |

### Saccade-rate

To test the prediction of the temporal orienting and the certainty hypotheses, we performed a two-way repeated measures ANOVA on the pre-target SR, with factors Condition (full-certainty, high-certainty-frequent, high-certainty-rare and low-certainty) and Foreperiod (1 s, 2 s). This analysis revealed a significant interaction between Condition and Foreperiod (*F*(3, 57) = 6.27,*p* < .001, 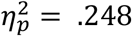, 95% CI = [.056 .407]), consistent with the temporal orienting hypothesis (**Figure 1B**). Main effects were not tested as there were no predictions regarding them and the interaction rendered them meaningless.

Next, to interpret the interaction and test the predictions of the temporal orienting and the certainty hypotheses, we analyzed the two foreperiods separately.

#### Analysis of 1 s foreperiod trials

We performed a one-way repeated-measures ANOVA on the pre-target SR of the 1 s foreperiod trials, with Condition as an independent variable. Consistent with both hypotheses, this analysis revealed a significant difference in mean pre-target SR across conditions (*F*(3, 57) = 13.36,*p* < .001, 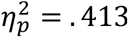, 95% CI = [.204 .556], **Figure 3A**). A planned contrast revealed a significant predictability effect: lower pre-target SR for blocks with a fixed foreperiod (full-certainty) relative to varying foreperiods (low-certainty) (*t*(57) = 5.442, FDR-adjusted *p* < .001, Cohen’s *d* = 1.721, 95% CI = [1.006, 2.436]). This effect, also consistent with both hypotheses, replicates our findings of a previous study [2].

Next, we performed two independent planned contrasts to test the predictions of the hypotheses: (a) high-certainty-rare vs. high-certainty-frequent; and (b) high-certainty-rare vs. low-certainty. The certainty hypothesis predicts that pre-target SR would be lower for high certainty relative to low certainty trials, but that it would be unaffected by foreperiod frequency. Therefore, it predicts that pre-target SR would: (a) not differ significantly between the high-certainty-rare and the high-certainty-frequent conditions; and (b) be lower for the high-certainty-rare relative to the low-certainty condition (**Figure 1A**). In contrast, the temporal orienting hypothesis predicts that target frequency would determine the pre-target SR and that certainty would have no effect. Therefore, it predicts that pre-target SR would: (a) be lower for the high-certainty-frequent relative to the high-certainty-rare condition; and (b) not differ significantly between the high-certainty-rare and the low-certainty conditions (as in both conditions frequency is 20%) (**Figure 1B**).

**Figure 3.**
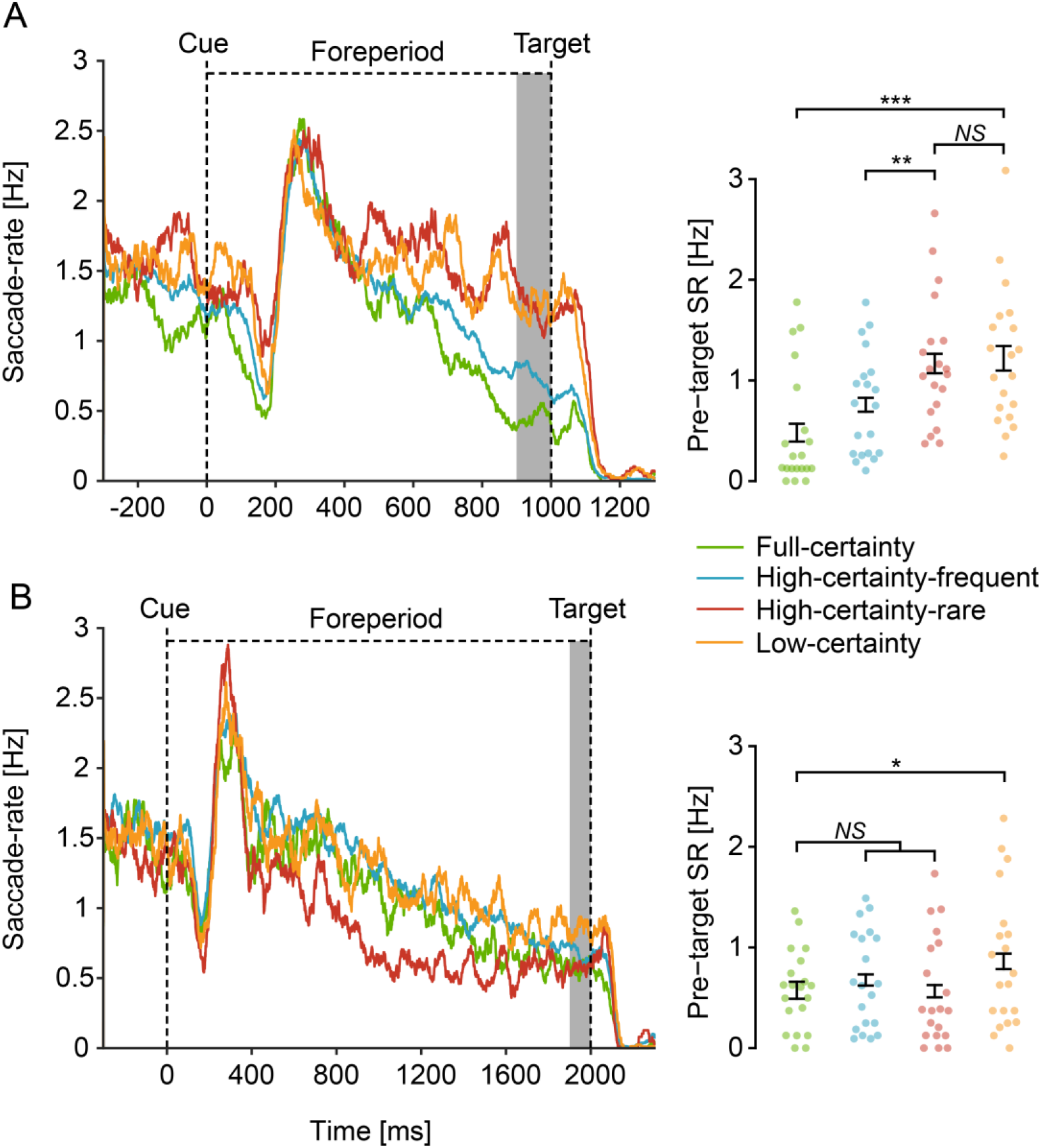
Saccade-rate results. Saccade-rate (SR) traces (left) relative to cue onset in 1 s (A) and 2 s (B) foreperiod trials, averaged across participants (N=20). SR traces were smoothed using 50 ms running window for display purposes. Mean pre-target SR was averaged across −100 to 0 ms relative to target onset (shaded region) and analyzed (right). Individual mean pre-target SR represented as dots for each condition. Error bars designate ±1 within-subject standard error from the mean [31]. FDR-corrected *p*-values are reported for the planned contrasts conducted. * *p* < .05; ** *p* < .01; *** *p* < .001; *NS* not significant.

Results showed that: (a) pre-target SR was lower for the high-certainty-frequent relative to the high-certainty-rare condition (*t*(57) = 3.015, FDR-adjusted *p* = .006, Cohen’s *d* = 0.953, 95% CI = [0.291 1.620]); and (b) there was no evidence for difference between the high-certainty-rare and low-certainty conditions (*t*(57) = 0.383, FDR-adjusted *p* = .703, *BF*_01_ = 4.13 ± .02%). This is consistent with the prediction of the temporal orienting hypothesis and inconsistent with the certainty hypothesis.

#### Analysis of 2 s foreperiod trials

We performed a one-way repeated-measures ANOVA on the pre-target SR of the 2 s foreperiod trials, with Condition as an independent variable. Consistent with both hypotheses, this analysis revealed a significant difference in mean pre-target SR across conditions (*F*(3, 57) = 3.798, *p* = .0149, 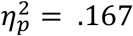, 95% CI = [.007 .32], **Figure 3B**). Using a planned contrast we found a significant predictability effect: lower pre-target SR for blocks with fixed foreperiods (full-certainty) relative to varying foreperiods (low-certainty) (*t*(57) = 2.876, FDR-adjusted *p* = 0.011, Cohen’s *d* = 0.909, 95% CI = [0.244 1.575]). This effect is only partially consistent with our previous study, where we found a predictability trend for trials of 2 s foreperiod, but this trend has not reached significance [2]. Interestingly, however, when reanalyzing the 2 s foreperiod trials from the previous study and matching the number of trials to the present study by including only the first 80 trials (as was done for the power analysis procedure, see Methods), the predictability effect was found to be significant also in the previous study.

To examine the alternative predictions for trials with a 2 s foreperiod, we contrasted the full-certainty condition against the two high-certainty conditions combined. The certainty hypothesis predicts that pre-target SR would be affected by certainty within a block rather than by the conditional probabilities. Therefore, it predicts lower pre-target SR for 2 s foreperiod trials in the full-certainty condition relative to the combined high-certainty condition (**Figure 1A**). In contrast, the temporal orienting hypothesis predicts that pre-target SR would be affected solely by the probability of the target to occur at a specific time, rather than by the general sense of certainty. Since the conditional probability for a target to occur at 2 s post cue reaches a maximum shortly prior to 2 s in both full-certainty and the two high-certainty conditions, this hypothesis predicts no difference in pre-target SR between these conditions. (**Figure 1B**)

There was no evidence for a significant difference in mean pre-target SR between the full-certainty and the combined two high-certainty conditions (*t*(57) = 0.541, FDR-adjusted *p* = .591, *BF*_01_ = 3.89 ± .02%). As in the 1 s foreperiod results, these results are consistent with the prediction of the temporal orienting hypothesis and inconsistent with the certainty hypothesis.

## Discussion

This study examined two competing explanations for the pre-target oculomotor inhibition effect. Under the temporal orienting hypothesis, pre-target oculomotor inhibition depends on the probability that the target would occur at a specific time. In contrast, under the certainty hypothesis, oculomotor inhibition depends on the participant’s state of certainty, which is determined by the consistency of target timings within a given context (i.e. a block). Results from the current study consistently support the predictions made by the temporal orienting hypothesis, and not those made by the certainty hypothesis.

In a series of previous studies it has been suggested that pre-target oculomotor inhibition could be an informative measurement for assessing target predictability [2–6]. These findings were interpreted as reflecting a link between pre-target oculomotor inhibition and three different types of temporal structures [1]: those due to the association between cue and target (associative regularity), those due to rhythmic stimulations (rhythmic regularity), and those due to conditional probabilities (hazard rate). Moreover, the pre-target oculomotor inhibition effect was found to be a more sensitive and a more reliable index for predictability than other common behavioral and electrophysical measurements [2].

The present study is an important step in strengthening this conclusion. The present findings provide a replication for the previously-reported effects [2–6]. As in previous studies, we found that: (a) slightly prior to target onset, saccade rate is lower (i.e. more strongly inhibited) for predictable relative to unpredictable targets; and (b) that for unpredictable targets, longer foreperiods are associated with lower pre-target saccade rates. The first finding supports the link between pre-target oculomotor inhibition and associative and rhythmic regularities, and the second finding supports the link to the hazard rate (further discussed in the Supplementary Material). In addition to providing replication, this study sheds new light on the interpretation of the present and previous findings by showing that the oculomotor inhibition effect does not reflect the general cognitive state of the participant, but can be directly linked to the anticipation of specific targets at specific timings.

Our definition of certainty in this study focuses on a long-term state of the participant which is assumed to be relatively constant within a block and not modulated by trials. In contrast, attentional orienting is a feature of each trial and is, by definition, a dynamic process that changes with time, and relative to stimulus presentation. It could have, therefore, been suggested finding a trial-based modulation by itself could disprove the certainty hypothesis. However, this is not the case. Trial-based modulations of saccade rate were found in all conditions including when foreperiods were random and targets were unpredictable. The onset of the cue, like any other visual stimulus, induces an inhibition of saccade rate, which is followed by a rebound and then by a gradual decrease until saccade rate reaches baseline [2]. This decrease until baseline could be long, lasting well into the foreperiod, and is likely the result of sensory or motor processes. Specifically it was previously suggested that alertness is increased with cue onset and this could be the result of the modulation in saccade rate in all conditions [21]. Therefore, finding such a trial-based modulation was not a sufficient support for the temporal orienting hypothesis, and it was necessary to compare experimental conditions as we did in this study.

Temporal expectation can be studied using a between-blocks or within-block design. In a between-blocks design (e.g. [2,22–25]), expectation is manipulated by comparing blocks with different foreperiod distributions; often fixed and random distributions. In a within-block design (e.g. [26–30]), expectation regarding target onset is manipulated by pairing symbolic cues with various target timings and by comparing trials with informative, uninformative and invalid cues.

Both between-blocks and within-block designs are valid and common approaches for studying temporal expectation, with each having its own advantages and disadvantages. Within-block designs eliminate most of the context-dependent effects and effects that slowly develop between blocks, such as fatigue and training. However, unlike between-block designs, the formation of expectations in within-block designs depends on a high-level identification of the semantic identities of multiple cues. Such high-level cue identification requires either explicit instructions or very long sessions to allow enough time to establish implicit learning of this complex information. In contrast, between-blocks designs, such as the one used in the current study, allow temporal regularities to be implicitly and quickly learned with no need for explicit instructions regarding the meaning of the cues.

Isolating the effects of temporal orienting from those of certainty, requires including either blocks (in a between-blocks design) or trials (in a within-block design) that manipulate these two factors separately. In a between-block design, this would require modifying the amount of predictable trials per block as was done in this study and previous ones (e.g. [24,25]). In a within-block design this would mean adding invalid trials, i.e. rare trials where the cue does not predict target onset (as in [29,30]). Importantly, while switching to a within-block design with invalid trials could have been a relatively straight-forward way to dissociate temporal orienting and certainty, doing so would have defeated the main goal of previous studies on oculomotor inhibition – to examine how oculomotor behavior reflects *implicit* learning of temporal regularities. By using a between-block design with a varied level of certainty, we demonstrated in this study how temporal orienting and certainty can be manipulated in a between-block design without compromising the implicit nature of the task. This approach could be advantageous for future studies in this field.

## Conclusions

This study shows that oculomotor inhibition reflects temporal orienting of attention. These findings support the feasibility of using pre-target oculomotor inhibition as a biomarker to study target-specific temporal expectation.

## Acknowledgements

This study was funded by the Israel Science Foundation grant 1960/19 to S.Y-G. We thank Omer Solomon for his assistance in the eye tracking analysis, and Marisa Ourieff for proof reading.

## Author Contribution

NTP and SYG designed this research and wrote the manuscript. NTP performed the experiment and analyzed the data.

## Data availability

The dataset used for power analysis, the dataset generated by this study, and an R-markdown file that reproduces all the reported statistical analyses, tables and bar graphs within the paper and supplementary material, are uploaded to the Open Science Foundation repository and are available at: https://osf.io/xyfsq/

