## Supplementary S1 for "Pre-target oculomotor inhibition reflects temporal orienting rather than certainty"

### *S1: Foreperiod effects - the involvement of conditional probabilities*

*Background*

Behavioral studies showed that when foreperiods vary within blocks, trials with longer foreperiods were associated with faster responses [1–3]. This behavioral *foreperiod effect* is thought to reflect changes in expectations due to the gradual increase in the conditional probabilities – the probabilities that a target would occur at a certain time given that it has not yet occurred [3]. Conditional probabilities follow a specific function called the 'hazard-rate' and therefore it is expected that neural and behavioral effects that are related to conditional probabilities would follow the same function. In a previous study, we found an oculomotor foreperiod effect – in blocks with varying foreperiod durations, longer foreperiods were associated with less pre-target SR, i.e. stronger oculomotor inhibition [4]. However, the interpretation of this effect remained inconclusive, as the change rate of the oculomotor inhibition effect across time was linear and did not closely follow a 'hazard rate' function.

*Results*

A one-way repeated measures ANOVA on the pre-target SR of low-certainty trials with foreperiod duration (1s/1.5s/2s/2.5s/3s) as the independent measure, revealed a significant difference between foreperiods ($F(4, 76) = 4.445$, $p = .003$, $\eta_p^2 = .19$, 95% CI = [.03 .32]); **Figure S1**). Polynomial contrasts revealed a significant negative linear trend

($t(76) = -3.810$, $p < .001$, Cohen's $d = -2.69$, 95% CI = [-1.2 -4.19]) with no additional significant polynomials ($p = .08 - .94$).

To assess the involvement of conditional probabilities in determining the pre-target SR, we calculated two Pearson correlations for each participant and condition: (a) between the pre-target SR (mean across trials) and the hazard-rate probability weights (0.2, 0.25, 0.33, 0.5, 1); and (b) between the pre-target SR and linear weights (0.2, 0.4, 0.6, 0.8, 1). Correlations were calculated individually for each participant and then transformed using Fisher Z-transformation using the psych package for R [5]. Mean correlations were calculated as the inverse Fisher transform of the averaged individual z-scores [6]. This analysis revealed that the pre-target SR significantly correlated with both a hazard-rate trend (mean $r_{hazard}$ across participants = -.28, $t(19) = -2.38$, $p = .028$) and a linear trend (mean $r_{linear}$ across participants = -.43, difference from zero: $t(19) = -3.13$, $p = .006$), but the correlation was stronger for the linear trend ($t(19) = -2.50, p = .022$).

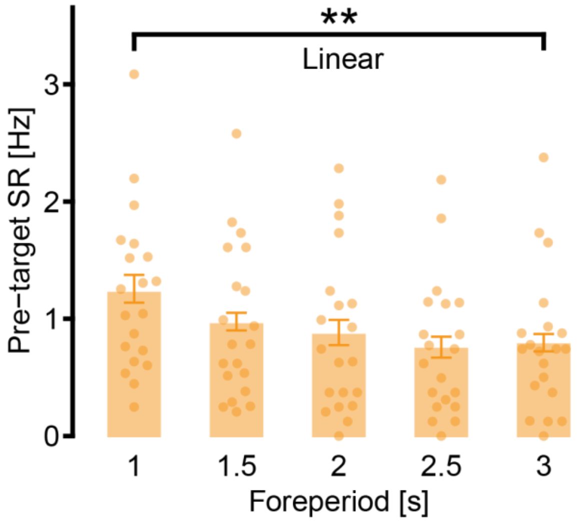


**Figure S1**. *Low-certainty trials results*. Mean pre-target saccade-rate (SR) averaged across -100 to 0 ms relative to target onset for each foreperiod, averaged across participants (N=20). Individual mean pre-target SR represented as dots for each foreperiod. Error bars designate ±1 within-subject standard error from the mean [11]. *p*-value indicated for polynomial contrasts. ** $p < .01$

*Discussion*

Findings of our previous study [4] provided inconclusive evidence regarding the link between pre-target oculomotor inhibition and “hazard-rate” expectations – expectations that are based on conditional probabilities. On one hand, our findings in the previous study [4], replicated here as well, showed that when the foreperiod varies within a block, oculomotor inhibition gradually increase as time progresses, (i.e. pre-target saccade rate was lower for a longer foreperiod). On the other hand, in both the previous [4] and current study, this effect fitted a linear trend better than an exponential hazard-rate modulation, which reflects changes in conditional probabilities as time progressed [4]. This poor fit with the hazard-rate function is consistent with previous findings on RTs using a similar design [1] and raises question regarding the interpretation that these effects are related to changes in expectations that are due to changes in conditional probabilities. As we have noted in our previous study, these findings are based on correlations across only five data points (the five foreperiods that were used). With only five data points, the distinction between the linear model and the hazard rate model is only minor and, therefore, any difference between them is difficult to interpret.

These findings are consistent with some studies that challenged the reliance of the perceptual system on the hazard-rate function for temporal expectation. The computation of the hazard-function over non-discrete time is relatively complex and numerically unstable [7,8]. A recent study found that the reciprocal of the probability density function (i.e., 1/PDF), which is simpler to compute than the hazard-rate function, provided a better fit to RT results in three separate sensory modalities [2]. Others have argued that the hazard-rate hypothesis fails to account for some of the known phenomena of temporal expectation [9]. It was further suggested that the commonly observed behavioral foreperiod effects can be explained by the degraded accumulation of memory traces from previous trials experienced by the participant, rather than by conditional probabilities [9,10]. Given the similarity between RT effects and the oculomotor inhibition effect observed here and in other studies [4], we acknowledge that these alternative accounts may also explain the pattern of results that we have observed without incurring the hazard-rate function. Importantly, this possibility does not affect the validity of the oculomotor inhibition as a marker of target-specific temporal expectation. Rather, converging evidence from this marker along with other markers may help resolve the open questions regarding the hazard-rate function.
